# Hormonal plasticity to food restriction is heritable

**DOI:** 10.1101/2023.09.29.559362

**Authors:** Jenny Q Ouyang, Ádám Z Lendvai

## Abstract

Theoretical and empirical studies agree that populations harbor extensive among-individual variation in phenotypic plasticity, but the mechanisms generating and maintaining this variation are often unknown. Endocrine systems that exhibit plastic changes in response to environmental variation may be subject to natural selection, but their evolution requires heritable variation. It is currently unknown if endocrine plasticity to environmental challenges is heritable. We tested whether glucocorticoid responsiveness to food restriction is heritable in house sparrows, *Passer domesticus*, by subjecting individuals to a standardized dietary restriction and selecting individuals according to their hormonal responsiveness to the treatments: into high plastic, low plastic, and control groups and let them reproduce. Using a cross-foster design, we compared the parental and the F1 generation to partition the heritability of glucocorticoid responsiveness into genetic and environmental sources of variation. We found moderate heritability (h^2^>20%) of glucocorticoid plasticity in response to food availability in both restricted and adequate food conditions. Environmental variance played a larger role under restricted than adequate food conditions, whereas residual variance was much higher under adequate food conditions. Our findings provide empirical evidence for the existence of heritable individual variation in glucocorticoid plasticity that selection can act upon, especially in rapidly changing environments.

## Introduction

A long-standing goal in evolutionary biology is to understand the relationship between environmentally-induced variation observed within a generation and genetically-based evolutionary changes between generations (West-Eberhard 1989). Although it is widely recognized that trait expression is plastic, the relationship between trait plasticity and its evolution remains an unresolved problem (Pigliucci 2005). Traditionally, environmentally-induced plasticity was viewed as non-heritable variation (West-Eberhard 2005; López-Maury et al. 2008). Currently, theoretical models recognize that environments can cause predictable patterns of plasticity that may enable organisms to rapidly adapt and survive in changing conditions (De Jong 2005; Ghalambor et al. 2007). Recent studies have documented a large degree of individual variation in plasticity (Nussey et al. 2005; Guindre-Parker 2020) that may interfere with the speed of evolutionary change and thus adaptation (Ghalambor et al. 2015). Despite its importance, the degree and persistence of individual reaction norms, *i*.*e*., pattern of phenotypic expression of a single genotype across a range of environments, responding to environmental variation remain less clear.

Theoretical and empirical studies agree that selection on plasticity can result in phenotypic adaptation to different environmental conditions, but the source of this plasticity can be a result of phenotypic or genetic variation and developmental conditions (Gavrilets and Scheiner 1993; DeWitt et al. 1998; Taff and Vitousek 2016). Theoretical models suggest that variation among individuals in plasticity may arise if there is spatiotemporal heterogeneity in resources resulting in frequency-dependent payoffs of alternative strategies (Wolf et al. 2008). Moreover, positive feedback mechanisms may reduce the costs of plasticity; therefore, stable coexistence of plastic and non-plastic individuals may arise due to heterogeneity in their previous experiences. Alternatively, stable individual differences in plasticity may arise due to genetic differences between the individuals, or these effects may even act in concert (Dingemanse and Wolf 2013). In order to discriminate between these possibilities, disentangling the additive genetic and the environmental effects are necessary (Araya-Ajoy and Dingemanse 2017).

Adaptive plasticity enhances the probability of population persistence in new environments and can facilitate genetic differentiation when directional selection acts on extreme phenotypes (Price et al. 2003). When the genotype interacts with the environment (G x E), different genotypes may respond differently to this environmental variation, resulting in variable fitness consequences when genotypes are expressed in different environments (West-Eberhard 1989). Natural selection can act on phenotypic variation in plasticity, but heritable variation is required for evolution to take place. Therefore, disentangling the genetic basis, the environmental context, and the interaction between genetics and environmental variation is critical for understanding phenotypic variation and organismal evolution.

Labile traits allow organisms to match their phenotype to environmental factors that vary within their lifetimes (Dingemanse et al. 2004). The key to responding adaptively to energetic demands and challenges is a system that integrates external and internal conditions and transmits that information to mediate resource allocation (Wingfield 2013). The hypothalamic-pituitary-adrenal (HPA) axis in vertebrates is such a system, releasing hormonal messengers, *e*.*g*., glucocorticoids (GCs), which regulate physiology and behavior in response to environmental challenges (Hau et al. 2016). At baseline levels, GCs regulate energy metabolism, *e*.*g*., gluconeogenesis (Exton 1979). Changes in GC concentrations show large individual plasticity (Taff et al. 2018; Houslay et al. 2019), and the degree of this plasticity has been shown to be associated with fitness parameters in the wild (Ouyang et al. 2013; Patterson et al. 2014; Sonnweber et al. 2018; Guindre-Parker et al. 2019). Therefore, activation of the HPA-axis represents a prime example of plastic reaction to environmental change. An important yet open question in evolutionary biology is whether endocrine plasticity is heritable, as the answer represents the first step towards understanding trait evolution and the functional significance of physiological plasticity.

When determining the heritability of a trait, we are estimating the contribution of genetic effects to the total phenotypic variance observed in that trait within a population (Pearson and Henrici 1896). Any change in the influence of the environment upon that trait will alter estimates of heritability, even when there is no change in the underlying genetic variation. Furthermore, phenotypic traits previously shaped by natural selection may lack standing genetic variation that would allow ongoing evolution. The level of plasticity varies among individuals, from non-responders to responders of varying degrees (Nussey et al. 2007; Brommer et al. 2008). A suite of studies in behavioral ecology has shown that this variation may be heritable (Dingemanse et al. 2012; Dochtermann et al. 2015; Araya-Ajoy and Dingemanse 2017), implying the evolution of plasticity. Endocrine plasticity, on the other hand, has received much less attention in evolutionary biology. Individuals have been shown to differ in their glucocorticoid plasticity (Lendvai et al. 2014; Sonnweber et al. 2018; Guindre-Parker et al. 2019; Houslay et al. 2019), showing remarkable intra- and inter-individual variation in response to varying environmental conditions (Vitousek et al. 2019; Zimmer et al. 2020). Recent work has shown that the slope and intercept of GC response can be repeatable under different environmental challenges (Baldan et al. 2021). While absolute glucocorticoid concentrations have been shown to be repeatable and heritable (Jenkins et al. 2014; Stedman et al. 2017; Bairos-Novak et al. 2018; Béziers et al. 2019), there is little evidence in the existence of heritable endocrine plasticity. Understanding the link between intra- and inter-generational variation is the first step to predicting organismal responses to environmental change and how selection may act on plasticity.

We use an environmentally relevant stressor of food restriction to test whether plasticity in glucocorticoid concentrations is heritable. We repeatedly sampled individual house sparrows, *Passer domesticus*, under restricted and control food conditions and assessed their initial glucocorticoid levels as a response to environmental variability. We then selected the most responsive, the least responsive and a random sample as a control group to produce a cross-fostered F1 generation and tested the F1 individuals’ glucocorticoid response to food restriction. Our main goal was to assess the environmental, genetic and residual contributions to glucocorticoid plasticity. We also assessed reproductive parameters to see if reproductive success is related to glucocorticoid plasticity.

## Methods

93 free-living juvenile house sparrows were caught in Reno, NV, USA over 3 days in fall 2016. After capture, birds were individually housed indoors under 12:12 LD in 47 × 31 × 36 cm cages. Birds were acclimated in indoor conditions for 30 days with *ad libitum* water and food. After 30 days, we measured each individual’s daily food consumption by offering 150g of millet mixed with chicken starter mix. For five consecutive days, 24 h after replenishing the individual food containers we removed husks and weighed the remaining food (including any spillage in the cage), and the difference between this value and the initial 150g was the daily food consumption (range: 6.8g-20.1g, some intake variation accounted for by sex and body weight). Average daily food consumption was calculated as the mean of the five-day food consumption values. As weight and volume of the seed mixture are highly correlated (Lendvai et al. 2014), to facilitate the distribution of daily food portions, we created individual-based volumes for 110% and 70% of the average individual daily food consumption. Treatment began after 5 days with two alternating weeks of food restricted (70%) and two control conditions (110%). After each week, we took a blood sample (∼70µl) from the brachial vein from each bird within 3 minutes of entering the room.

### Selection

After the dietary treatment was concluded, we determined corticosterone values (see below) from the blood samples collected after each treatment. We rank ordered the 93 individuals based on their responsiveness (steepness of the slope of the corticosterone levels over food restricted and control conditions calculated as the slope of the line of best fit through the points in a linear model). We chose 60 individuals (20 most reactive (10 males and 10 females), 20 least reactive (10 males and 10 females), and 20 (10 males and 10 females) randomly selected from the remaining 53) (Figure 1) and separated these groups of individuals into 3 large outdoor flight aviaries (5m x 5m). We note that the 20 most reactive individuals were not perfectly balanced by sex but that we chose the 10 most reactive females and 10 most reactive males (likewise for least reactive).

**Figure 1.**
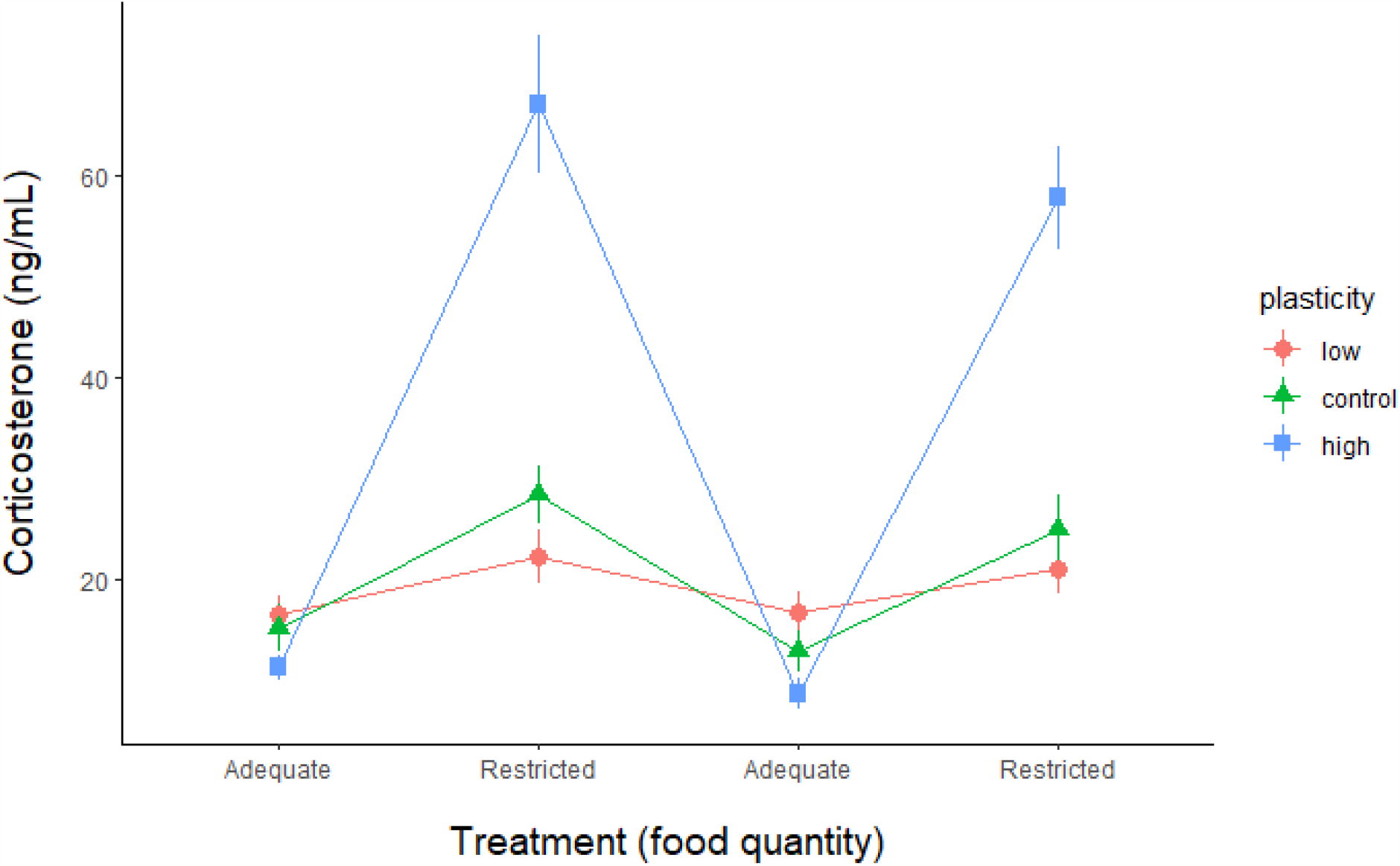
Corticosterone responsiveness (means ± SE) in the three experimental group selected for different levels of endocrine plasticity. House sparrows (n=20 per group) selected for low (red, circle), high (blue, square), and control (green, triangle) corticosterone plasticity in response to food restriction treatment formed the parental generation.

This parental generation was given *ad libitum* food and water refreshed daily, supplemented with dried mealworms and fresh fruit once a week. We provided nest boxes over winter and nesting material and mealworm pupae early spring (March) to stimulate breeding. House sparrows line up their nest with feathers shortly before egg laying, therefore we provided feathers at the same time to all aviaries in late March to stimulate similar breeding phenology. Sparrows paired in March and started the first clutches in April (all three aviaries within 10 days of each other). We checked nest boxes daily for presence of eggs and incubation start. Once incubation started, boxes were not checked again until the expected date of hatching to minimize disturbance. Once the young hatched, we marked individuals with a combination of nail clipping and nail polish and fostered half of the offspring to a synchronous nest in the same aviary within 2 days of the last young hatched. We chose individuals randomly but tried to match the cross-fostered young with those of similar weight. We performed the crossfoster within each aviary rather than among aviaries, as we are interested in maintaining selection lines of phenotypic plasticity. Original clutch sizes were always maintained during cross-fostering. Once the first nestlings hatched, we provided each aviary with ad lib live mealworms, frozen crickets, and wet cat food for parents to feed their offspring. Once young were independent (∼90 days after fledging), we transferred them into adjacent outdoor aviaries. In the following fall, the F1 generation (n=78; juveniles) were all tested for corticosterone response to food treatment in the same way as their parents (and at the same period). Post-fledgling mortality was high (although not as high as in the wild) due to competition with adult birds (fledglings were often chased), limited post-fledgling care (adults did not feed), and attempted predation and injury from a pair of hawks. For reproductive parameters, we used hatch date as the first date eggs hatched, clutch size as the number of eggs within a clutch, hatching success as the number of eggs that hatched out of the total number of eggs in a clutch, and fledgling number as the number of young to successfully leave the nest.

Extrapair paternity (EPP) occurrences in our aviaries are too low to account for in our analyses (0.04 ± 0.02 proportion of extrapair chicks/brood). Briefly, we isolated genomic DNA from both nestlings and parents from a subset of the population and carried out a polymerase chain reaction to amplify five microsatellite regions (Neumann and Wetton 1996; Griffith et al. 1999; Stewart et al. 2006). We calculated critical values such that the tolerance of mismatches was set to accept mismatches up to 2.73 (95%) and 1.09 (85%). We categorized an individual as extrapair if there was one or more mismatches and CERVUS-based analyses did not recognize the social father as the most likely father (Ouyang et al. 2014). These EPP rates are lower than free-living populations of house sparrows, but they are comparable to food-supplemented house sparrows, likely due to the fact that aviary males can more easily and continuously monitor their mates during laying (Václav et al. 2003).

### Hormone assays

To measure plasma corticosterone, we used enzyme-linked immunosorbent assay kits (Enzo Life Sciences; Farmingdale, NY, USA) following the manufacturer’s instructions, with a standard curve on each plate. Based on previous validations of this assay (Baldan et al. 2021), we determined that house sparrow plasma should be diluted 1:40 with 0.5% steroid displacement reagent. We randomly assigned samples in triplicates across plates, with the exception that all samples from the same individual were on the same plate. We included a standard curve on each plate, which ranged from 32 pg/mL to 20,000 pg/mL. The assay sensitivity was 2.1 pg/ml. To calculate intra- and inter-plate coefficient of variation (CV), we also included three pooled house sparrow samples on each plate, and each pool was assayed in triplicate. The intra-plate CV was 6.8% and inter-plate CV was 4.3%.

### Statistical analyses

Data processing and statistical analyses were carried out in the R computing environment using R version 3.6.3 (R Core Team). First, we used a univariate repeated-measures model using Bayesian Markov Chain Monte Carlo (MCMC) estimation implemented in the R package MCMCglmm (Hadfield 2010). The goal of this model was to analyze the effects of the dietary treatment on the circulating corticosterone levels in both the parental and the F1 generation and across the plasticity groups. In this model, we analyzed log-transformed corticosterone values measured after the dietary treatments as the dependent variable. Fixed effects included the food treatment (adequate: 110% and restricted: 70% of average daily food intake), age (parental or F1 generation), the three plasticity groups (factor with three levels: low or high plasticity group and controls) and all possible interactions of these three factors.

In the next step, we ran a random regression animal model on the log-transformed corticosterone values. The model was fitted using corticosterone values as dependent variable and the two dietary treatments as fixed factors. The adequate treatment can be interpreted as the intercept, and the restricted treatment as the responsiveness (i.e. plasticity) of the individuals to the food shortage. Total phenotypic variance (VP) was partitioned into the following components: among-individual variance, additive genetic effects (VA), common environmental effects (VE) and residual variance (VR). From these components, we calculated the narrow-sense heritability (h^2^ as VA/VP) and the effect of common rearing environment (e^2^ as VE/VP). VA and VE variance components were estimated by including pedigree information and the identity of the foster nest (rearing environment) into the model as random factors. Contributing factors to residual variation (VR) can include unaccounted variation in environmental conditions, internal state, and biological instabilities, or simple measurement error. Fixed effects were not included into this model to avoid ambiguity regarding the estimates of heritability. Intercept and slope heritabilities were estimated as the amount of additive genetic variance in intercepts and slopes extracted from the animal model divided by their respective total phenotypic variance (*i*.*e*., the sum of the additive genetic, permanent environment and among-treatment variances). An identical statistical approach has been used to analyze the heritability of plasticity for behavioral traits (Via et al. 1995; Araya-Ajoy and Dingemanse 2017). We had no information about the genetic relatedness of the parental generation; therefore, all adults were considered to be unrelated. In all models, we used an inverse Wishart prior (a weakly informative prior with minimal influence on the posterior distribution). Models were run for 500.000 iterations, with 10.000 burn-in and a thinning interval of 200. We report posterior means and 95% credible intervals (CI) and Bayesian p-values. For illustration purposes, in the figures (Fig. 2-3) hormonal responsiveness is shown as the slope from a linear model fit through the log corticosterone values measured under restricted and adequate food conditions. Convergence of the models were estimated using a combination of visual and numerical approaches. For visual verification, we checked the trace plots and appropriate mixing of different chains. Numerically, the convergence was tested using the Gelman-Rubin and Geweke’s criterion using the ‘gelman.diag’ and ‘geweke.diag’ and ‘geweke.plot’ functions in the ‘coda’ package (Plummer et al. 2006). Reproductive parameters were also analyzed in univariate MCMCglmm models, with nest included as a random factor in the models.

**Figure 2.**
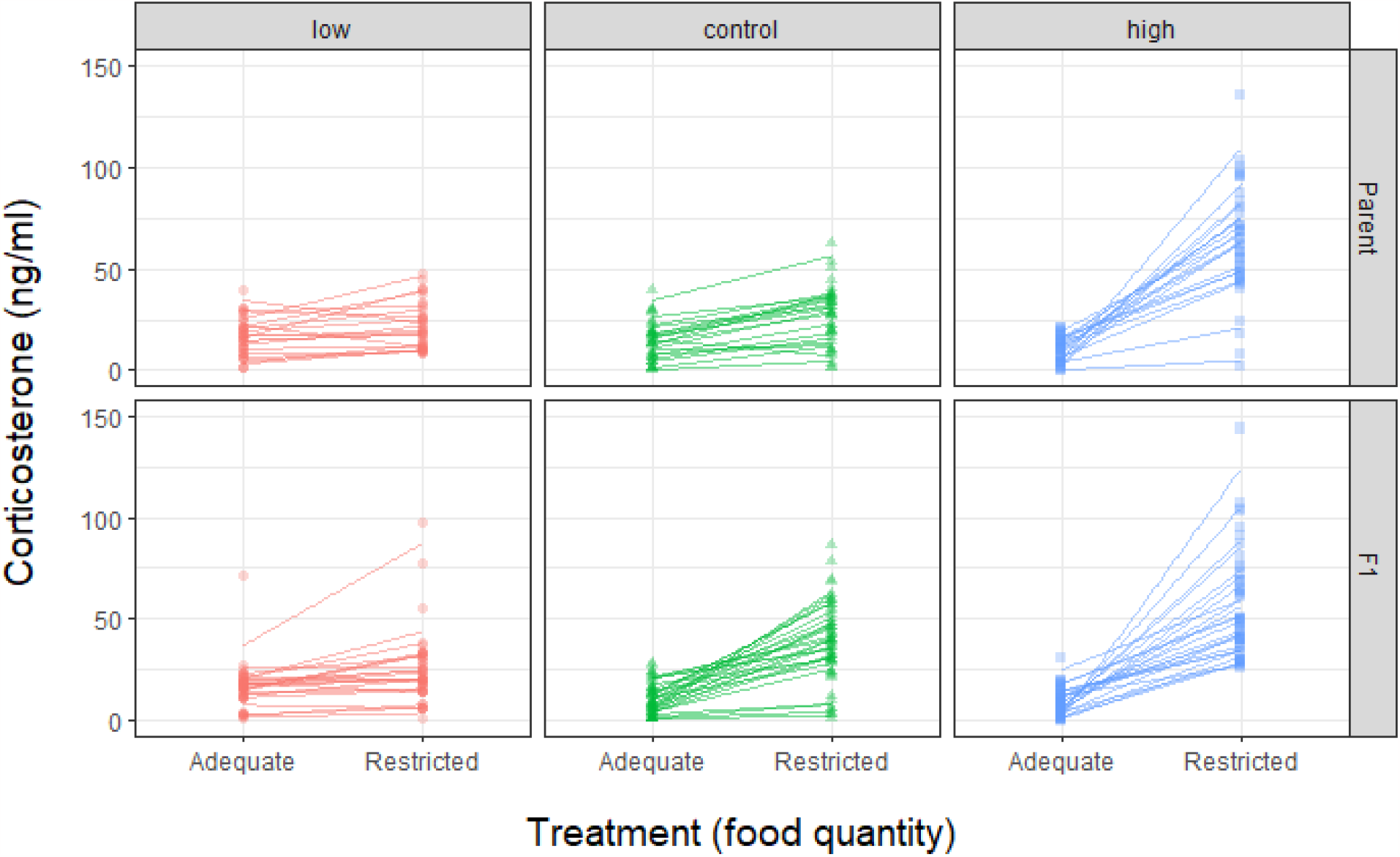
Hormonal response to food restriction in the parental and the F1 generation of house sparrows in three selection lines. The columns show the selection lines (low, control and high plasticity groups), and the rows show the parental and the F1 generation.

## Results

### Treatment effectiveness and difference between selection lines

Under the ‘adequate food’ treatment, corticosterone concentrations did not differ between the parental and F1 generations or between plasticity groups (Table S1). However, food restriction induced an increase in corticosterone levels (0.88 [0.58;1.19], p < 0.001). Compared to the control group, corticosterone responsiveness was higher in the high plasticity group (1.27 [0.86;1.72], p<0.001), and lower in the low plasticity group (-0.52 [-0.92;-0.06], p = 0.017). In the control group, individuals in the F1 generation had stronger responsiveness than their parents (generation × food treatment interaction: 0.58 [0.14;0.96], p = 0.004). However, compared to the controls, responsiveness tended to increase less in the F1 generation in high and low plasticity individuals (Fig. 2), resulting in similar values to their parents (generation × food treatment × plasticity interaction; high plasticity: -0.58 [-1.16;-0.04] p = 0.039, low plasticity: -0.55[-1.12;0.01], p = 0.064, Fig. 2, see Table S1 for full model results). In juveniles, the hormonal response differed between the plasticity groups (Table S2a), but was not different between male and female birds (Table S2b). Adding paternal or maternal influence to these models did not increase model fit (Table S2c-d; Fig. S1).

### Heritability

Variation in corticosterone levels were heritable (Fig. 3). When we quantified the proportion of heritable variance (Table 1, Table S3) we found that the hormonal response to the challenging restricted food condition (i.e. reaction norm slope) had similar heritability (20%) than the hormone levels measured at adequate food conditions (i.e. reaction norm intercept) (21%). The effect of common rearing environment (e^2^) was low for the reaction norm intercept but was stronger for the reaction norm slope (Table 1). Among-individual variance also explained a larger part for the responsiveness (slope) than for the levels measured under adequate food conditions (intercept). Within-individual (residual) variance was higher for the reaction norm intercept than for slope (Table 1).

**Table 1.** Proportion of the total phenotypic variance under the two experimental conditions. Posterior mode and in brackets 95% Bayesian credible intervals are shown.

| | $h^2$ (narrow sense heritability) | $e^2$ (environmental) | individual ID | R (residual) |
| --- | --- | --- | --- | --- |
| **Intercept** | 21.91 [9.65-36.79] | 11.18 [5.07-30.23] | 22.1 [9.18-35.11] | 37.24 [28.2-49.56] |
| **Responsiveness** | 20.32 [9.3-35.5] | 30.32 [12.24-49.51] | 36.52 [18.69-52.19] | 12.02 [7.81-16.8] |

**Figure 3.**
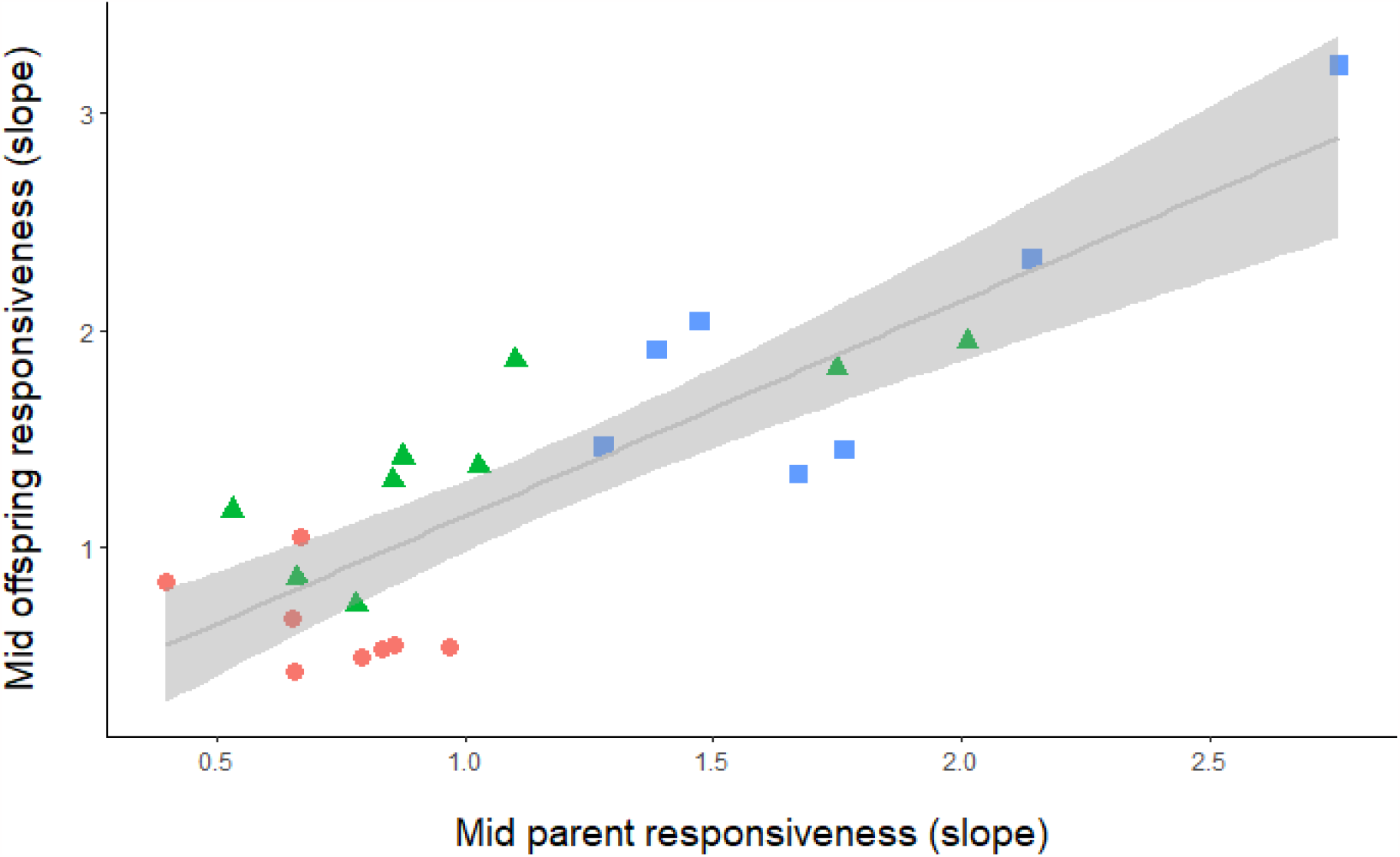
Mid-offspring and mid-parent correlation between the three groups. Red dots, green triangles and blue squares indicate the low, control and high plasticity groups, respectively. For illustration purposes only, corticosterone responsiveness was calculated as the difference between the average corticosterone levels under the two experimental dietary conditions (*i*.*e*., the slope on a log-scale, ng/ml).

### Reproductive parameters of parental generation

Overall, 80 % of adults in outdoor aviaries successfully bred and raised offspring to fledgling. The remaining 20% either did not pair up or in the case of 3 females, laid eggs but did not incubate. Birds in the high plasticity group were less likely to breed successfully (70%), than the low plasticity group (80%), while the most successful were the controls (90%) (χ^2^ = 10, df = 2, p = 0.006). However, among birds that did breed, high plasticity birds (along with controls) started breeding sooner (2.41[1.10;4.95] days sooner; p = 0.022, Fig. 4a, Table S4a). There was no statistical difference in clutch size, hatching success, and fledgling number between the plasticity groups (Figure 4b-d, Table S4b-d, p > 0.2).

**Figure 4.**
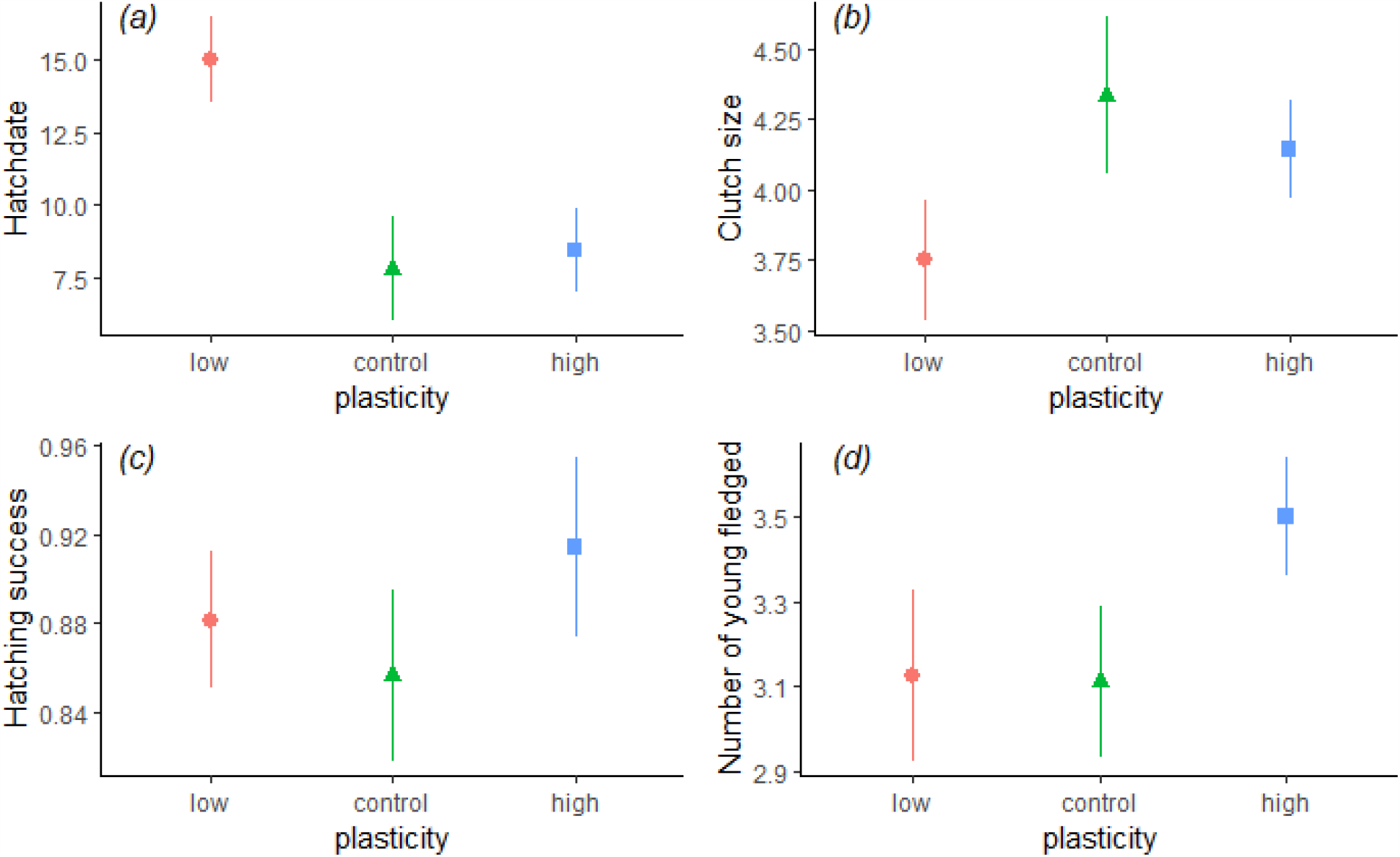
Reproductive parameters (means ± SE) in adult house sparrows differing in their endocrine plasticity. Adult sparrows showing different glucocorticoid plasticity in response to food restriction. Among birds that bred, low plasticity individuals started breeding significantly later (hatch date with 1= 1^st^ of April) than control and high plasticity individuals (a). Clutch size (b), hatching success (c), and fledgling number (d) did not differ significantly between the groups. See Supplementary materials (Table S4) for full model outputs.

## Discussion

We determined whether glucocorticoid plasticity is heritable using an environmentally relevant challenge, food restriction. We found moderate heritability (20-21%) of both initial corticosterone levels and their plasticity (*i*.*e*., responsiveness to food restriction).

Previously, we reported significant individual variation in hormonal responsiveness to changes in food availability in house sparrows (Lendvai et al. 2014). In this study, not only do we confirm this result, but we also show that this phenotypic variation is heritable, *i*.*e*., is due to additive genetic effects. Environmental variance seems to be especially important during challenging environmental conditions, corroborating earlier results (Jenkins et al. 2014). It is possible that when resources are limited, variance can be easily attributed to either environmental or genetic factors. However, when resources are sufficient, it is more difficult to measure phenotypic variance as environmental variance increases. We have good evidence both in field and laboratory conditions that absolute glucocorticoid concentrations are heritable (Wada et al. 2008; Jenkins et al. 2014; Béziers et al. 2019). Additionally, if glucocorticoid responsiveness is also heritable, the HPA system (either at the level of the hypothalamus, pituitary or the adrenals) is likely under strong selection via differential contribution to reproductive success, especially during early stages of divergence in new environments (Lande 2009). It is also during this stage that plasticity will either reduce or exacerbate initial mismatch between average and optimal phenotypic responses. We tested the glucocorticoid response to food restriction, which is likely related to physiological pathways related to metabolism and digestion. Our heritability estimates also come from controlled laboratory conditions, and we note that the dynamic nature of heritability in natural populations may be different, likely overestimated in laboratory conditions with reduced environmental sources of variation. It is necessary to assess the variation and heritability of other endocrine systems, possibly under other environmental variations, to gain more insight into endocrine evolution.

What can be the cause of such heritable variation in endocrine responsiveness? Early life conditions, starting from maternal investment to variation in rearing environments, are likely to cause differences in endocrine responsiveness (Monaghan 2008; Lendvai et al. 2009; Goerlich et al. 2012). For example, early nutritional stress in avian species leads to altered stress physiology in adulthood, from the molecular level to behavioral and physiological changes (Jimeno et al. 2017; Zito et al. 2017; Grace and Anderson 2018). In our study, cross-fostering had little effect on differences between the genetic and rearing environments, because nestlings were only cross-fostered within the treatment groups, *i*.*e*., between parents with similar phenotypic responses to food restriction. However, early maternal effects, such as glucocorticoid deposition altering yolk hormone levels, parental feeding behavior and sibling competition still differed between different rearing nests leaving still a potentially large source of environmental variation that could affect offspring phenotypes (Love and Williams 2008). A limitation to our study is the *ad libitum* food resources in our aviaries, which is unrealistic in nature and may limit our ability to detect differences between genetic and environmental rearing conditions.

The fact that we found significant and heritable among individual variation in glucocorticoid plasticity is important because it means that not only the average baseline hormone concentrations, but their responsiveness (reaction norm) can evolve under natural selection. Physiological plasticity can both constrain and facilitate evolution (Ghalambor et al. 2015; Diamond and Martin 2016). For example, if plasticity confers higher mean fitness, the variation in reaction norms in the population can weaken subsequent selection by hiding genotypic variation, because different genotypes may express the same phenotypic response under some environmental conditions. Maladaptive plasticity can even drive a population to extinction through homeostatic failure (Cotto et al. 2019). Both cases would result in evolutionary constraint. On the other hand, physiological plasticity can facilitate evolution by buffering populations from extirpation so that selection can act on standing, *i*.*e*., cryptic genetic variation (Ghalambor et al. 2007). An appropriate endocrine response in dynamic environments is likely required to optimize organismal fitness. The natural population from which we sampled contained both non-responsive and responsive individuals. The coexistence of both “phenotypes” suggests that an evolutionary mechanism maintains the natural variation in plasticity within populations. For example, if variation in environmental conditions (food availability, precipitation, etc.) among years is high, physiologically plastic individuals may do better in one year than the next (Luttbeg et al. 2021). Indeed, two recent studies found that adrenal sensitivity was highest in unpredictable environments (Zimmer et al. 2020; Guindre-Parker and Rubenstein 2021). Alternatively, if individuals do not often experience large variation in the environmental conditions, then the genotype by environment interaction (G × E) may mask the genetic differences among individuals. In this case, under stable conditions (which may not be unrealistic for house sparrows that are obligate commensalists of human settlements with stable conditions year-round), individuals with different genotypes may express similar phenotypes. These scenarios allow for both phenotypes to exist within changing environmental pressures.

Although our experiment was set out to quantify the heritability of glucocorticoid responsiveness, breeding in the three plasticity groups provided some insights to the adaptive value of different phenotypes, as they differed in some reproductive parameters. Since we did not have replicates within experimental groups, we acknowledge that the reproductive differences could be attributed to an ‘aviary effect’ (albeit unlikely, since all outdoor aviaries had similar enrichment, water and food availability, temperature and humidity). Although we provided identical breeding conditions, the high plasticity individuals were less likely to breed. One biological explanation for the propensity to breed could be related to the negative effect of high glucocorticoid levels on the hypothalamic pituitary gonadal axis. Highly plastic individuals exhibit higher circulating levels of corticosterone, possibly to additional stressors besides food availability, and this long-term effect of glucocorticoids impacts the HPG axis and may activate GnIH, directly inhibiting reproduction (Bentley et al. 2006). However, if these individuals bred, they started breeding earlier than control and less plastic pairs. Our birds are breeding in controlled aviary conditions with *ad libitum* access to food and water, perhaps representative of a predictable environment. In general, constant or predictable environments may allow for more bet hedging or proactive personalities, or those with less of a stress response (Cockrem 2007). Under unpredictable rainfall patterns, for example, baboons were more likely to have increased stress responses (Gesquiere et al. 2008). It could be that only the best quality individuals in the high plasticity group were able to, or chose, to breed. There is evidence that variation in exploratory behavior co-varies with corticosterone concentrations (Baugh et al. 2012), so it could be that more exploratory individuals that do well are also those that are more responsive physiologically to food restriction. We note that reproduction differences among three groups could also be driven by individual past experiences (age, developmental conditions, illness), which might affect life history traits as well as the HPA axis.

Understanding the evolution of labile phenotypic traits requires the analysis of relative contributions of genetic, environmental and developmental factors to assessing reaction norms. Here we show evidence of the existence of heritable individual variation in glucocorticoid plasticity, which is a first step into recognizing the different means by which plasticity can contribute to adaptive evolution. The next challenge is to analyze how this variation is related to fitness. Although differences in the reproductive parameters among our groups in differing plasticity is suggestive of a phenology differences, it requires independent experimental confirmation. Approaches such as selection or introduction experiments in nature, following populations over time, will provide an opportunity to measure patterns of plasticity and the rate of genetic differentiation.

## Notes

https://zenodo.org/record/8377792

### Competing Interest Statement

The authors have declared no competing interest.

