## Supplementary materials for "Hormonal plasticity to food restriction is heritable"

Supplemental materials for **Hormonal responsiveness to food restriction is heritable in a songbird**

**Results**

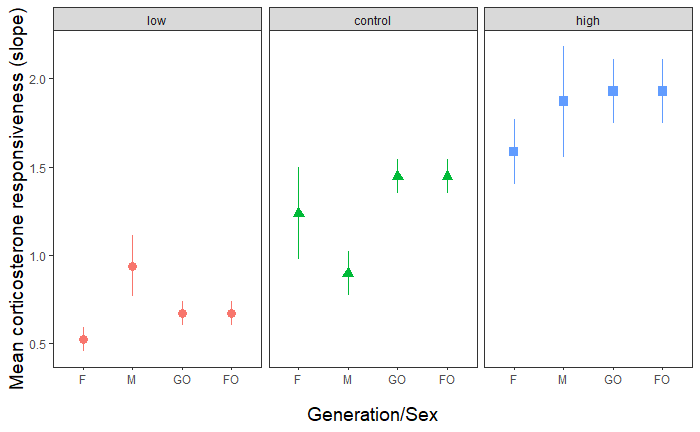

**Figure S1. Corticosterone responsiveness (mean ± SE) in the F1 individuals over the three selected groups: low plasticity, high plasticity, and control,** separated by generations (F = female, M = male parents, and GO and FO are offspring in the F1 generation) and whether individuals in the F1 generation were the foster (FO) or genetic offspring (GO). For illustration purposes only, corticosterone responsiveness was calculated as the difference between the average corticosterone levels under the two experimental dietary conditions (i.e., the slope on a log-scale, ng/ml). Corticosterone responsiveness in the F1 generation was higher than in their parents in the control group.

**Table S1. Full model results on corticosterone plasticity using a MCMC estimation analyzing the effects of dietary treatment on circulating corticosterone levels in both parental and F1 generations across plasticity groups.** Fixed effects include food treatment (treat, treatL= restricted 70% of intake, treatH= adequate 110% of intake), age (ageJuv=F1, ageAdult=parental), the three plasticity groups (low, high, control). The intercept corresponds to the 110% intake, parental generation and control group.

```
Location effects: log(cort) ~ treat * age * plasticity

(Intercept)                    2.18602  1.83835  2.52169     2450 < 4e-04 ***
treatL                         0.88545  0.58961  1.19355     2450 < 4e-04 ***
ageJuv                        -0.25186 -0.71432  0.20694     2878 0.28735
plasticitylow                  0.39764 -0.11076  0.87979     2599 0.12000
plasticityhigh                -0.36936 -0.89087  0.07459     2450 0.12571
treatL:ageJuv                  0.58645  0.14592  0.96868     2637 0.00408 **
treatL:plasticitylow          -0.52661 -0.92144 -0.06419     2320 0.01714 *
treatL:plasticityhigh          1.27447  0.86461  1.72717     2450 < 4e-04 ***
ageJuv:plasticitylow           0.24653 -0.43255  0.89098     2803 0.44735
ageJuv:plasticityhigh          0.17699 -0.46949  0.81330     2450 0.60163
treatL:ageJuv:plasticitylow   -0.54821 -1.12574  0.01351     2450 0.06449 .
treatL:ageJuv:plasticityhigh  -0.57741 -1.16038 -0.04553     2450 0.03918 *

---
Signif. codes:  0 ‘***’ 0.001 ‘**’ 0.01 ‘*’ 0.05 ‘.’ 0.1 ‘ ’ 1
```

**Table S2. Full model results on corticosterone plasticity using a MCMC estimation analyzing the effects of dietary treatment on circulating corticosterone levels in the F1 generations across plasticity groups.** Fixed effects include food treatment (treat, treatL= restricted 70% of intake, treatH= adequate 110% of intake), age (ageJuv=F1, ageAdult=parental), the three plasticity groups (low, high, control) and sex (M = males, F = females). The intercept corresponds to the 110% intake, parental generation, control group and females (in model (b) where sex is included).

**(a)**

```
DIC: 707.212

Location effects: log(cort) ~ treat * plasticity

                       post.mean   l-95% CI   u-95% CI eff.samp  pMCMC
(Intercept)           -4.057e+01 -2.494e+04  1.835e+04     2450  0.707
treatL                 1.470e+00  1.207e+00  1.709e+00     2450 <4e-04 ***
plasticitylow          2.453e+02 -3.212e+04  2.653e+04     2593  0.795
plasticityhigh        -1.327e+02 -2.502e+04  3.315e+04     2668  0.891
treatL:plasticitylow  -1.071e+00 -1.408e+00 -7.005e-01     2450 <4e-04 ***
treatL:plasticityhigh  7.014e-01  3.476e-01  1.076e+00     2266 <4e-04 ***
```

**(b) effect of sex in juveniles**

```
DIC = DIC: 706.7158

Location effects: log(cort) ~ treat * plasticity * sex

                            post.mean   l-95% CI   u-95% CI eff.samp  pMCMC
(Intercept)                 2.059e+02 -2.230e+04  2.459e+04     2450  0.659
treatL                      1.550e+00  1.191e+00  1.903e+00     2242 <4e-04 ***
plasticitylow               3.041e+02 -3.194e+04  2.748e+04     2450  0.810
plasticityhigh              4.107e+02 -3.206e+04  3.191e+04     2450  0.848
sexM                       -1.026e-01 -6.973e-01  4.097e-01     2450  0.740
treatL:plasticitylow       -1.155e+00 -1.655e+00 -6.709e-01     2450 <4e-04 ***
treatL:plasticityhigh       8.848e-01  4.009e-01  1.391e+00     2450 <4e-04 ***
treatL:sexM                -1.530e-01 -6.557e-01  3.635e-01     2243  0.536
plasticitylow:sexM          4.541e-01 -3.034e-01  1.306e+00     2450  0.268
plasticityhigh:sexM         6.140e-01 -1.592e-01  1.473e+00     2450  0.130
treatL:plasticitylow:sexM   1.717e-01 -5.631e-01  9.116e-01     2450  0.666
treatL:plasticityhigh:sexM -5.085e-01 -1.262e+00  2.179e-01     2450  0.177
---
Signif. codes:  0 ‘***’ 0.001 ‘**’ 0.01 ‘*’ 0.05 ‘.’ 0.1 ‘ ’ 1
```

**(c) including maternal effect**

```
DIC = 707.2396


Location effects: log(cort) ~ treat * plasticity

                       post.mean   l-95% CI   u-95% CI eff.samp  pMCMC
(Intercept)            1.971e+01 -1.832e+04  1.882e+04     2450  0.652
treatL                 1.468e+00  1.214e+00  1.715e+00     2450 <4e-04 ***
plasticitylow          3.603e+01 -3.030e+04  2.049e+04     2450  0.791
plasticityhigh        -1.209e+02 -2.886e+04  2.162e+04     2450  0.885
treatL:plasticitylow  -1.066e+00 -1.457e+00 -7.176e-01     2450 <4e-04 ***
treatL:plasticityhigh  6.977e-01  3.488e-01  1.085e+00     2450 <4e-04 ***
---
Signif. codes:  0 ‘***’ 0.001 ‘**’ 0.01 ‘*’ 0.05 ‘.’ 0.1 ‘ ’ 1
```

**(d) including paternal effect**

```
DIC: 707.212


Location effects: log(cort) ~ treat * plasticity

                       post.mean   l-95% CI   u-95% CI eff.samp  pMCMC
(Intercept)            1.276e+02 -1.931e+04  2.118e+04     2971  0.667
treatL                 1.471e+00  1.224e+00  1.732e+00     2450 <4e-04 ***
plasticitylow          4.087e+02 -3.235e+04  2.766e+04     2450  0.776
plasticityhigh         4.753e+01 -2.883e+04  2.907e+04     3033  0.920
treatL:plasticitylow  -1.070e+00 -1.462e+00 -7.073e-01     2450 <4e-04 ***
treatL:plasticityhigh  6.953e-01  3.266e-01  1.052e+00     2450 <4e-04 ***
---
Signif. codes:  0 ‘***’ 0.001 ‘**’ 0.01 ‘*’ 0.05 ‘.’ 0.1 ‘ ’ 1
```

**Table S3. Full model results of MCMCglmm animal model and raw variance-covariance matrix effects**

```
Location effects: log(cort) ~ treat

            post.mean l-95% CI u-95% CI eff.samp  pMCMC
(Intercept)    2.1659   1.9447   2.4009     2384 <4e-04 ***
treatL         1.1820   0.9284   1.4318     2602 <4e-04 ***
---
Signif. codes:  0 ‘***’ 0.001 ‘**’ 0.01 ‘*’ 0.05 ‘.’ 0.1 ‘ ’ 1
```

```
treatH:treatH.animal     treatL:treatH.animal     treatH:treatL.animal     treatL:treatL.animal
0.30105826              -0.01312013              -0.01312013               0.15923459
treatH:treatH.band       treatL:treatH.band       treatH:treatL.band       treatL:treatL.band
0.32232599               0.20293307               0.20293307               0.29220845
treatH:treatH.fosternest treatL:treatH.fosternest treatH:treatL.fosternest treatL:treatL.fosternest
0.21018584               0.03787637               0.03787637               0.19777075
treatH:treatH.units      treatL:treatH.units      treatH:treatL.units      treatL:treatL.units
0.64084584              -0.02593013              -0.02593013               0.10682277
```

**Table S4. Model results for reproductive parameters**

(a) hatching date

```
Location effects: hatchdate ~ plasticity

                  post.mean l-95% CI u-95% CI eff.samp  pMCMC
(Intercept)          2.6004   2.1148   3.0776    649.4 <4e-04 ***
plasticitycontrol   -0.8878  -1.6033  -0.2492    796.1 0.0142 *
plasticityhigh      -0.5968  -1.3150   0.1090    546.3 0.0969 .
---
Signif. codes:  0 ‘***’ 0.001 ‘**’ 0.01 ‘*’ 0.05 ‘.’ 0.1 ‘ ’ 1
```

(b) Clutch size

```
Location effects: clutch ~ plasticity

                  post.mean l-95% CI u-95% CI eff.samp  pMCMC
(Intercept)          3.7588   2.9948   4.4518    417.4 <1e-04 ***
plasticitycontrol    0.5880  -0.3809   1.5940    673.9  0.234
plasticityhigh       0.4163  -0.5856   1.4254    420.2  0.409
---
Signif. codes:  0 ‘***’ 0.001 ‘**’ 0.01 ‘*’ 0.05 ‘.’ 0.1 ‘ ’ 1
```

(c) Hatching success

```
Location effects: cbind(hatch, I(clutch - hatch)) ~ plasticity

                  post.mean l-95% CI u-95% CI eff.samp   pMCMC
(Intercept)          2.3913   1.0286   4.0701    677.5 0.00384 **
plasticitycontrol   -0.2039  -2.0370   1.6909   1902.4 0.79636
plasticityhigh       0.5807  -1.4895   2.8536    929.7 0.56343
```

(d) Number of fledglings:

```
Location effects: fledge ~ plasticity

                  post.mean l-95% CI u-95% CI eff.samp  pMCMC
(Intercept)         1.05723  0.79544  1.27176    57.62 <2e-05 ***
plasticitycontrol   0.07587 -0.23115  0.40294    88.93  0.507
plasticityhigh      0.16140 -0.18935  0.38782    92.08  0.234
---
Signif. codes:  0 ‘***’ 0.001 ‘**’ 0.01 ‘*’ 0.05 ‘.’ 0.1 ‘ ’ 1
```
